# Dynamic patterns of YAP1 expression and cellular localization in the developing and injured utricle

**DOI:** 10.1101/2020.05.06.081323

**Authors:** Vikrant Borse, Matthew Barton, Harry Arndt, Tejbeer Kaur, Mark E. Warchol

## Abstract

The Hippo pathway is an evolutionarily conserved signaling pathway involved in regulating organ size, development, homeostasis and regeneration^1–4^. YAP1 is a transcriptional coactivator and the primary effector of Hippo signaling. Upstream activation of the Hippo pathway leads to nuclear translocation of YAP1, which then evokes changes in gene expression and cell cycle entry^5^. A prior study has demonstrated nuclear translocation of YAP1 in the supporting cells of the developing utricle^6^, but the possible role of YAP1 in hair cell regeneration is unclear. The present study characterizes the cellular localization of YAP1 in the utricles of mice and chicks, both under normal conditions and after hair cell injury. During neonatal development of the mouse utricle, YAP1 expression was observed in the cytoplasm of supporting cells, and was also transiently expressed in the cytoplasm of hair cells. We also observed temporary nuclear translocation of YAP1 in supporting cells of the mouse utricle at short time periods after placement in organotypic culture. However, little or no nuclear translocation of YAP1 was observed after injury to the utricles of neonatal or mature mice. In contrast, a significant degree of YAP1 nuclear translocation was observed in the chicken utricle after streptomycin-induced hair cell damage *in vitro* and *in vivo*. Together, these data suggest that differences in YAP1 signaling may be partly responsible for the distinct regenerative abilities of the avian vs. mammalian inner ear.

## Introduction

The sensory hair cells of the inner ear convert mechanical stimuli into electrical signals that mediate the senses of hearing and balance. Ongoing interactions between hair cells, their neighboring supporting cells and afferent and efferent neurons is essential for proper sensory function^7^. In mammals, loss of hair cells leads to permanent deficits in hearing and equilibrium^8–9^ In contrast, the ears of nonmammals can regenerate new hair cells after acoustic trauma or ototoxic injury^10–13^. The vestibular organs of mammals possess a limited capability to produce new hair cells^14^, but the extent of regeneration may not be sufficient for complete recovery of sensory function.

At the cellular level, two distinct mechanism has been shown to produce new hair cells after damage^15^. In some sensory organs, supporting cells can re-enter the cell cycle in response to hair cell injury, resulting in the production of new hair cells and supporting cells^11–13^. In other cases, supporting cells can undergo direct trans-differentiation, converting into new hair cells without proliferating^16–18^. Multiple signaling pathways, such as Notch, Wnt, FGF and VEGF, have been shown to be involved in hair cell regeneration and functional recovery after damage^19–20^. Hair cell-specific transcription factors and other transcriptional regulators such as p27^Kip1^, GATA3, ATOH1, and POU4F3 are also involved in the regenerative process^21–24^. Present knowledge of the pathways responsible for regeneration are incomplete, but their identification will be essential for development of methods for restoration of function in the inner ears of humans.

The Hippo/YAP1 pathway is an evolutionarily conserved signaling network known to be involved in regulating tissue size and cell number during development^2–4,25–26^, The transcriptional coactivator YAP1 is the primary effector of Hippo signaling. Under normal conditions, YAP1 is sequestered in the cytoplasm and targeted for degradation. However, activation of upstream Hippo pathway molecules or mechanical stimulation of cells and tissues can result in the nuclear translocation of YAP1, leading to changes in gene expression that promote cell division^5^. YAP1 signaling has been shown to play an important role in the development of the mouse utricle. Gnedeva et al., (2017) report that reduced mechanical stress in the sensory epithelium of the growing utricle promotes nuclear translocation of YAP1 and increased proliferation. It is not clear, however, whether YAP1 signaling is also required for regeneration after hair cell injury.

The present study profiled the expression pattern of YAP1 in the neonatal and mature mouse utricle and investigated the role of YAP1 signaling after selective hair cell lesion. We found that, during neonatal development, YAP1 is present in supporting cells, and is also transiently expressed in hair cells. We also observed transient nuclear translocation of YAP1 in supporting cells of mouse utricles shortly after placement in organotypic culture. However, selective hair cell ablation in Pou4f3-huDTR mice did not induce significant YAP1 nuclear translocation. In contrast, hair cell injury caused nuclear translocation of YAP1 in the chicken utricle. The results suggest that YAP1 can respond to mechanical forces acting on the sensory epithelium, but that hair cell injury in the mammalian utricle is not sufficient to promote YAP1 entry into the nucleus. Our data also reveal differences in the regulation of regeneration in the avian vs. mammalian inner ear.

## Results

### Developmental profile of YAP1 expression in the mouse utricle *in vivo*

Initial studies characterized the expression pattern of YAP1 in the mouse utricle during the first 15 days of postnatal development. Previous data indicate that growth and differentiation in the mouse utricle initially occurs in the striolar and medial regions, with the lateral region being the last to differentiate^27–28^. We obtained images from the lateral extrastriolar (L), striolar (S) and medial extrastriolar (M) regions of whole mount utricles at postnatal days P0, P7 and P15. Specimens were immunolabeled for myosin Vlla (hair cells), Sox2 (supporting cell and type II hair cells) and YAP1. Consistent with earlier studies^6^, we found that YAP1 expression was mainly confined to the cytoplasm of supporting cells and that its expression levels decreased with time. Interestingly, we also observed YAP1 expression in the cytoplasm of a subset of hair cells until P7, but YAP1 expression in hair cells was lost by P15 (Fig. 1A and 1B). Predominantly, YAP1-expressing hair cells were also immunoreactive for Sox2 and were most numerous in the lateral region of the sensory epithelium. Quantitative analysis verified that the numbers of YAP1^+^/Sox2^+^ hair cells diminished during postnatal development (Fig. 1C). In agreement with earlier findings^27–28^, we also observed a significant increase in Sox2^−^ hair cells and an increase in hair cell density between P0 and P15 (Fig. 1D).

**Figure 1.**
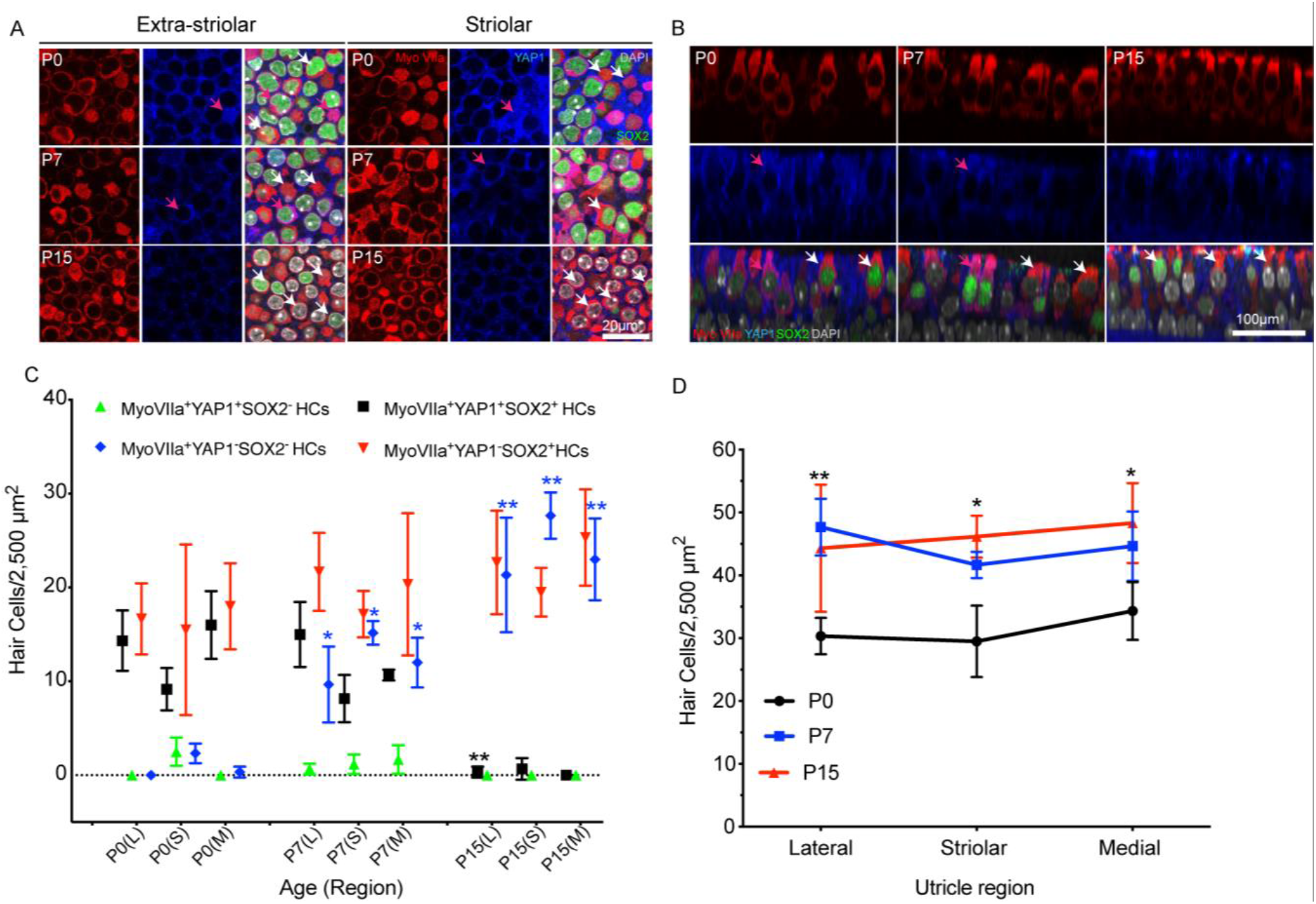
YAP1 expression profile in the developing mouse utricle *in vivo*. Mouse utricles were collected at postnatal days 0 (P0), P7 and P15. Immunostaining was performed using antibodies against Myosin VIIa (hair cell marker, red), SOX2 (green), YAP1 (blue). All cell nuclei were labeled with DAPI (grey). One or two images were taken from lateral and medial extrastriolar regions and from striolar region of each utricle. (A) YAP1-positive hair cells (magenta) were observed at P0 and P7, but were very rare at P15. Magenta arrows indicate MyoVlla^+^YAP1^+^ hair cells (magenta) and white arrows indicate MyoVlla^+^YAP1^-^ hair cells (red). Representative images were taken from XY planes of Z-stack images. (B) Representative orthogonal images showing YZ plane of P0, P7 and P15 mouse utricles. Magenta arrows indicate MyoVlla^+^YAP1^+^ SOX2^+ /-^ hair cells (magenta) and white arrows indicate MyoVlla^+^YAP1^−^ SOX2^+/-^ hair cells (red). (C, D) Quantitative data on utricle hair cell density, YAP1 and SOX2 expression during postnatal development. All data were obtained from 50 X 50 μm regions within the lateral extrastriolar, striolar and medial extrastriolar regions of each utricle. (C) During postnatal development, we observed significant increases in MyoVlla^+^YAP1^−^SOX2^−^ hair cells (blue) and decreases in MyoVlla^+^YAP1^+^SOX2^+^ hair cells (black), relative to P0. MyoVlla^+^YAP1^-^SOX2^+^ hair cells (red) and MyoVlla^+^YAP1^+^SOX2^−^ hair cells (green) did not change (relative to P0). (D) Increases in hair cell density were observed throughout the utricle at P15, relative to P0. Data expressed as mean + SD. Statistical test was two-way ANOVA followed by Tukey’s post hoc test (*p indicate significance relative to P0 and **p indicate significance relative P0 as well as P7, p value < 0.05). Supporting cells bar 10 μm. N=3-6 utricles.

Utricles fixed at P0 and P3 also displayed evidence of cell division. Mitotic figures at the metaphasic and anaphasic stages were occasionally observed in Sox2-expressing cells in the lateral region of the utricle. Nuclei of these cells showed light granular YAP1 immunolabeling (Fig. 2A). Utricles fixed between P0 and P7 also contained a small number of ‘atypical’ hair cells, which possessed large globular cytoplasmic structures and slightly elongated nuclei. Such cells were observed in medial and striolar regions of the sensory epithelium (density: 1-2 per 10,000 μm^2^) and displayed cytoplasmic immunoreactivity for YAP1 (Fig. 2B). We further found that the size of hair cell nuclei underwent a significant decrease during postnatal development (Fig. 2D). At P0, the average hair cell nuclear area in the striolar region was 72.40 + 14.24 μm^2^. At P7 and P15, average hair cell nuclear area in the striola was decreased to 44.23 + 3.35 μm^2^, and 43.50 + 4.50 μm^2^, respectively. A similar pattern was observed in the extrastriolar regions (Fig. 2D). Regression correlation analysis between hair cell density and nuclear area indicated a strong negative (Pearson’s) correlation, with r value of −0.8377 and p value < 0.0001 (Fig. 2E).

**Figure 2.**
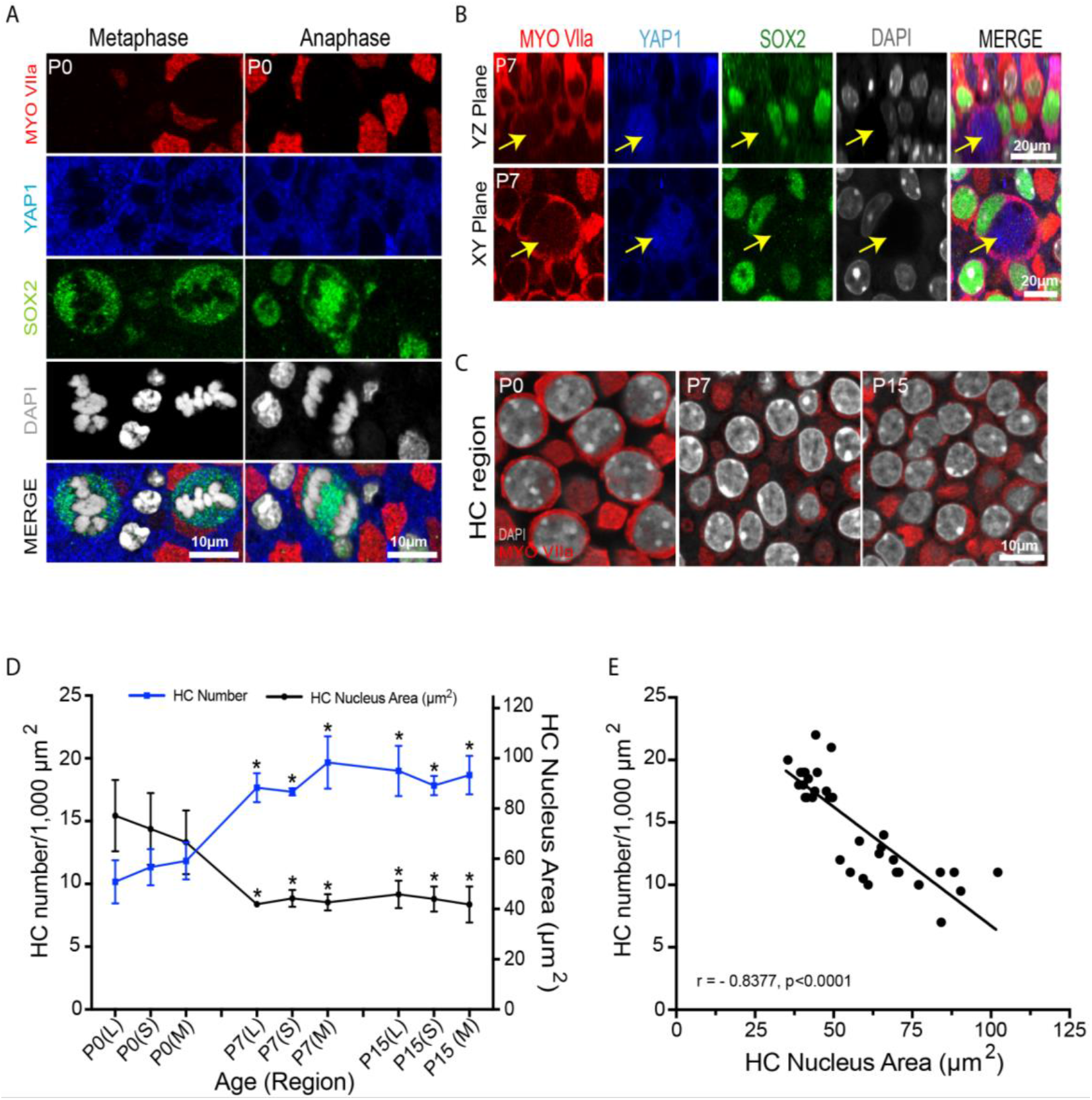
Mitotic cells, atypical hair cells and decrease in hair cell nuclear area in the developing mouse utricle. (A) Confocal images of mitotically dividing cells showing metaphasic and anaphasic chromosome alignment in whole mount utricle at P0, immuno label for Myosin VIIa (red), YAP1 (blue), SOX2 (green) and DAPI (grey). Light granular labeling of YAP1 was observed in mitotically dividing SOX2-positive cells. (B) High level of YAP1immunolabeling (blue) was observed in a small population of Myosin VIIa (red) and SOX2 (green) double-positive ‘atypical hair cells (yellow arrow).’ Images were obtained at P7. (C) Confocal images from hair cell region of the developing mouse utricle, showing changes in the size of hair cell nuclei. Labels: Myosin VIIa (red), and DAPI (grey). (D-E) Quantitative data on utricle hair cell density and hair cell nuclear area during postnatal development. All data were obtained from 1,000 μm^2^ regions within the lateral extrastriolar, striolar and medial extrastriolar regions. (D) At P7 and P15, a significant increase in hair cell number and decrease in hair cell nucleus area was observed throughout the utricle, relative to P0. (E) Significant negative Pearson correlation was observed between hair cell number and nuclear area in the developing mouse utricle. Data expressed as mean + SD. Statistical test used two-way ANOVA followed by Tukey’s post hoc test (*p value < 0.05 relative to P0). N=3-6 utricles.

### Placement in organotypic culture evokes transient nuclear translocation of YAP1

Nuclear translocation of YAP1 is regulated by multiple mechanical stimuli, such as matrix stiffness, epithelial stretching and cell density^2–4,25–26^. To test the effects of mechanical environment on YAP1 localization, we removed utricles from P15 mice and placed them in organotypic culture. Explanted utricles were placed in Matrigel-coated Mat-Tek dishes that contained 100 μl of culture medium. The surface of the sensory epithelium was placed in contact with the Matrigel substrate (i.e., lumenal side down) and a small amount of pressure was applied to ensure attachment. Cultured utricles were fixed after 2-24 hr *in vitro* and were then labeled with Sox2 and YAP1 antibodies. Examination of these specimens revealed transient nuclear translocation of YAP1 in supporting cells between 2-6 hr. *in vitro* (Fig.3A). After 2 hr in culture, nuclear YAP1 was observed in 68.20 + 15.39 % and 70.30 + 20.43% of supporting cells in the extrastriolar and striolar regions, respectively. After 6 hr, the degree of YAP1 nuclear translocation was observed in 75.14 + 15.63 % of extrastriolar and 79.49 + 12.44% of striolar supporting cells. In contrast, utricles that had been in culture for 24 hr contained 9.26 + 9.70 % and 17.79 + 14.69% YAP1-labeled nuclei in extrastriolar and striolar regions (respectively). Such data suggest that placement in culture leads to temporary YAP1 nuclear translocation, but that nuclear immunoreactivity of YAP1 returned to normal (0 hr time point) levels by 24 hr in vitro (Fig. 3C).

**Figure 3.**
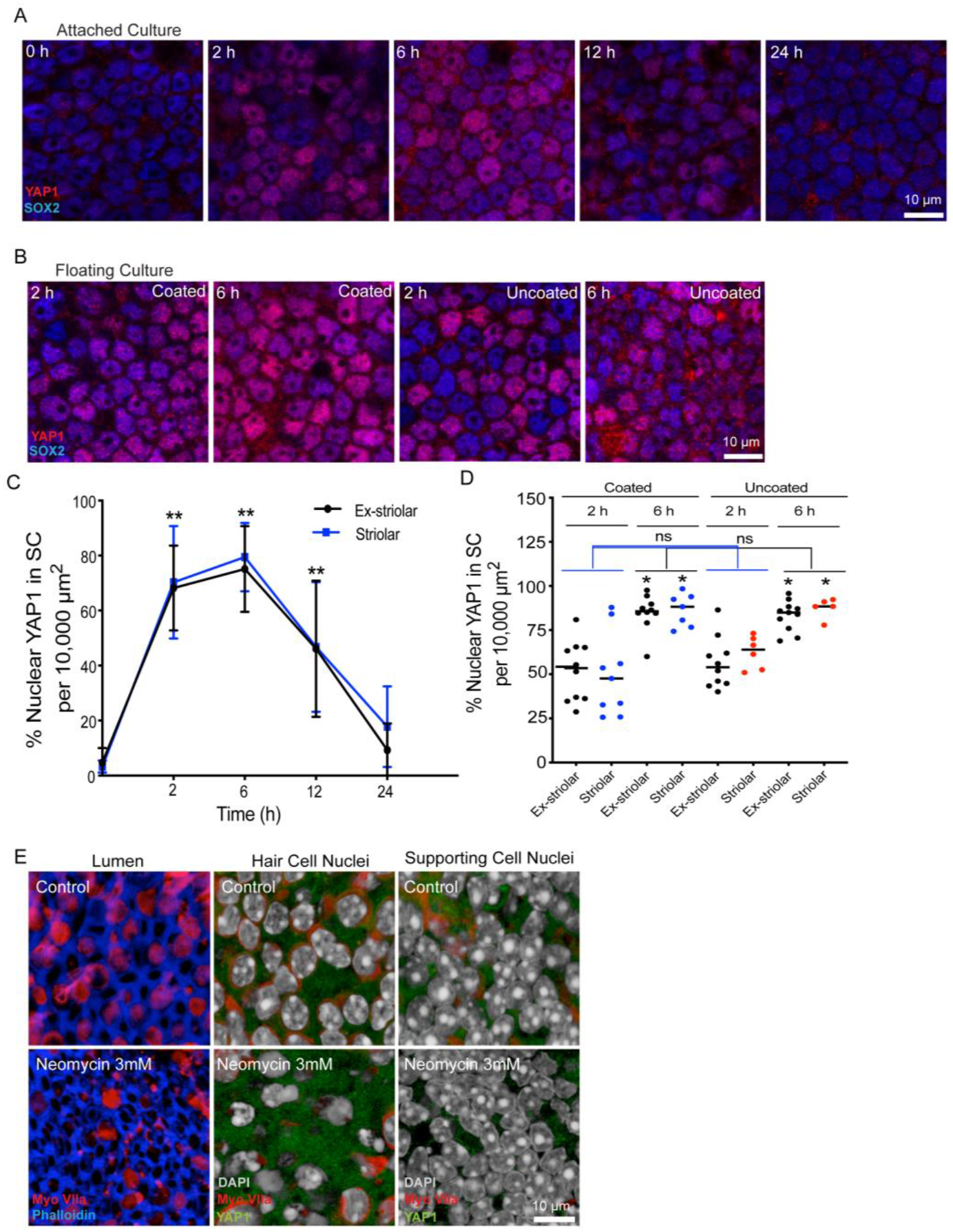
Transient nuclear translocation of YAP1 in organotypic culture of the mouse utricle. (A) Utricles were explanted from CD1 mice at P15, and attached to Matrigel-coated dishes. Specimens were fixed and examined after 0-24 hours in culture. Confocal images show supporting cells of cultured utricles immunolabeled for YAP1 (red) and SOX2 (blue). (B) Utricles were explanted at P15 and cultured in a Matrigel-coated or uncoated dishes as free-floating samples. Cultures were fixed and examined after 2 and 6 hours *in vitro*. Confocal images showing supporting cells of cultured utricles immunolabeled for YAP1 (red) and SOX2 (blue). (C) Quantitative data on the percentage of supporting cells with nuclear YAP1 immunoreactivity. All data were obtained from 10,000 μm^2^ regions within the extrastriolar and striolar regions of each utricle. There was a significant increase in nuclear YAP1 immunolabeling at 2hr, 6hr and 12hr *in vitro*, relative to specimens fixed immediately after explanation (0hr). (D) Quantitative data on the percentage of supporting cells with YAP1-labeled nuclei. All data were obtained from 10,000 μm^2^ regions within the extrastriolar and striolar regions of each utricle. There was no significant increase in the percentage of nuclear YAP1 at 2hr and 6hr time points between the coated and uncoated cultures. However, there was a significant increase in nuclear YAP1 immunolabeling in utricles cultured in uncoated dishes at the 6hr time point, vs. those cultured for 2hr. (E) Treatment of cultured utricles with neomycin resulted in loss of hair cells (left and middle columns), but did not lead to nuclear immunoreactivity for YAP1 in supporting cells (right column). Data expressed as mean + SD. Statistical test use one-way ANOVA followed by Bonferroni’s post hoc test (*p value < 0.05) (**p-value< 0.05 for Ex-striolar and Striolar both region). N=3-6 utricles.

The transient YAP1 nuclear translocation that was observed in cultured utricles may have been caused by attachment to the Matrigel coated dish. To test this, we cultured mouse utricles as free-floating samples in both Matrigel-coated and uncoated MatTek dishes. Utricles were maintained *in vitro* for 2 hr and 6 hr, since we previously observed maximum translocation at those times (Fig. 3). Following fixation and immunoprocessing, we observed a similar degree of YAP1 nuclear translocation in specimens cultured in both Matrigel-coated and uncoated dishes (Fig. 3B, D).

The patterns of YAP1 immunolocalization in the mouse utricle were also unaffected by ototoxic injury. Utricles were explanted from mice at ~1 month postnatal and placed in culture in uncoated MatTek dishes. Some utricles (n= 5) were treated with 3 mM neomycin, while control specimens (n= 5) were cultured identically, but without neomycin. After 24 hr in vitro, specimens were fixed and processed for labeling of hair cells and YAP1. Immunolabeling of myosin VIIa (Fig. 3E) indicated that neomycin treatment led to a reduction in hair density (17.6+8.8 hair cells/2,500 μm^2^ in neomycin treated utricles vs. 42.8+4.8 hair cells/2,500 μm^2^ in controls, p=4.5 x 10^−7^). However, very similar patterns of YAP1 localization were observed in both neomycin-treated and control utricles. Occasional YAP1-labeled nuclei were observed (data not shown), but they were rare and did not differ between lesioned utricles and controls.

### Selective hair cell ablation in Pou4f3-huDTR mice does not cause nuclear translocation of YAP1

Additional experiments examined epithelial repair and YAP1 localization following lesions to the mouse utricle. These studies used Pou4f3-huDTR transgenic mice, in which one allele of Pou4f3 is replaced by a gene encoding the human form of *HBEGF* (the ‘diphtheria toxin receptor’), which then facilitates the internalization of diphtheria toxin in hair cells^27^. Treatment of mature mice Pou4f3-huDTR mice with 25 ng/gm diphtheria toxin (DT) leads to partial loss of vestibular hair cells^18,28^. Pou4f3-huDTR and Pou4f3+/+ (WT control) mice (4-6 weeks of age) received a single 25 ng/ml injection of DT, and utricles were examined after 7- and 14-days recovery. Fixed specimens were immunolabeled for myosin Vlla, Sox2 and YAP1 and imaged using confocal microscopy. Resulting data showed loss of myosin VIIa-labeled hair cells in the striolar and extrastriolar regions of Pou4f3-huDTR mice, relative to WT controls. However, YAP1 nuclear translocation was not observed in any region of either the Pou4f3-huDTR or WT utricles at either time point (Fig. 4A).

**Figure 4.**
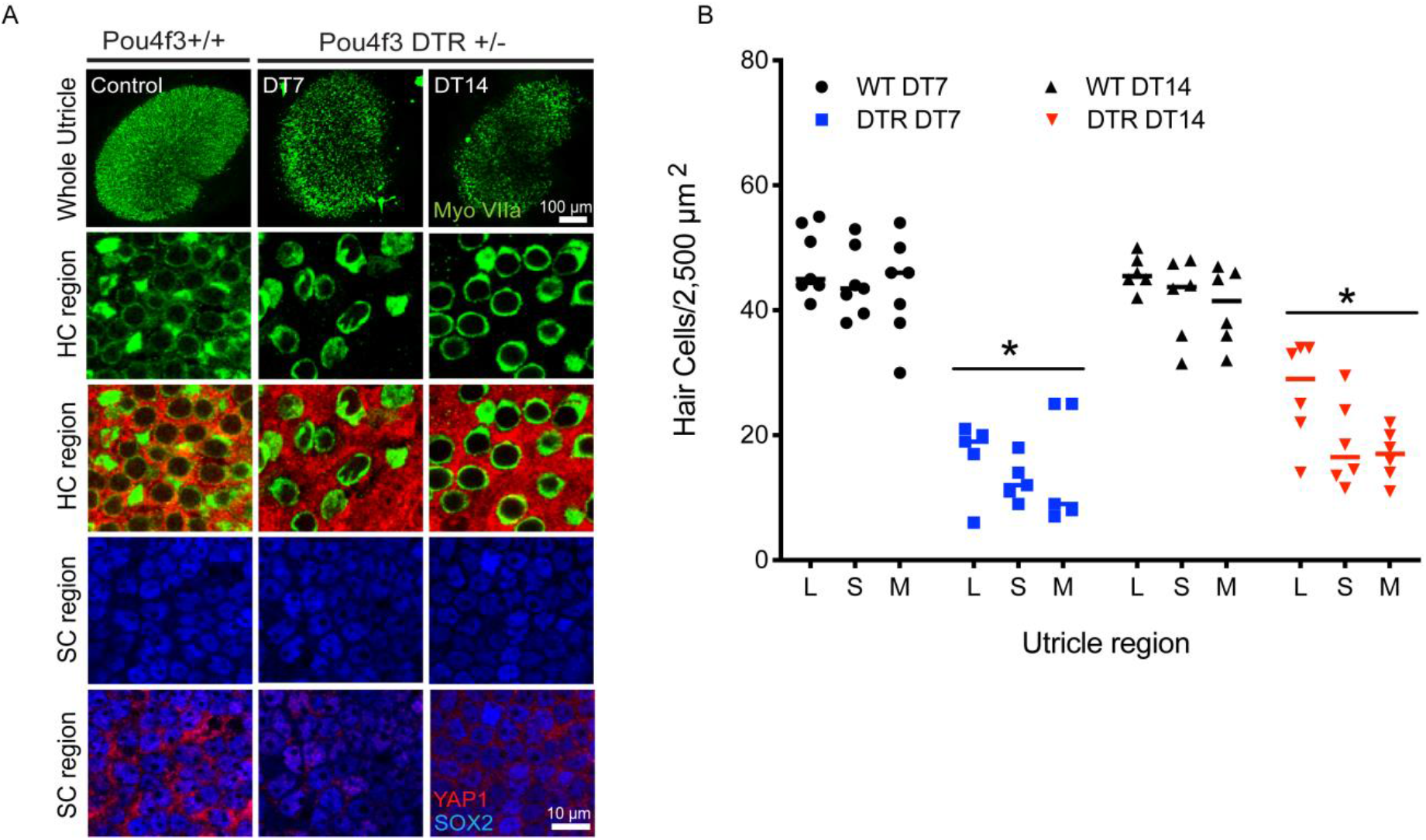
Nuclear translocation of YAP1 was not observed after diphtheria toxin (DT)-mediated hair cell lesions. (A) Adult Pou4f3 +/+ and Pou4f3 +/DTR mice were injected with a single dose of DT (25ng/g i.p). Utricles were collected, fixed and immunolabeled for myosin VIIa (green), SOX2 (blue) and YAP1 (red) after 7 days (DT7) and 14 days (DT14). Confocal images of wholemount utricles from Pou4f3 +/+ (wild type control) (DT14) and Pou4f3 +/DTR DT7 and DT14 are shown in the top panel. Images from the hair cell region and supporting cell region are shown in the second, third panels and fourth, fifth panels respectively. Treatment with DT led to reduced hair cell numbers (green) and increased area of YAP1 positive (red) labeling in Pou4f3 +/DTR at DT14 and DT7, as compared to wild type control. (B) Quantitative data on utricle hair cell density and YAP1 nuclear expression after a single dose of DT (25ng/g i.p). All data were obtained from 50 X 50 μm regions within the lateral extrastriolar, striolar and medial extrastriolar regions of each utricle. There was significant decrease in hair cell numbers at DT7 and DT14, compared to controls. However, this hair cell loss did not lead to YAP1 immunoreactivity in the nuclei of supporting cells. Data expressed as mean + SD. Statistical tests used two-way ANOVA followed by Tukey’s post hoc test (*p value < 0.05 relative to P0). N=4-6 utricle.

We further examined whether a severe lesion to the sensory epithelium was capable of causing nuclear translocation of YAP1. Unexpectedly, we found that a single 5 ng/gm injection of DT injection given to Pou4f3-huDTR mice at P5 resulted in a massive lesion in the sensory epithelium that involved the loss of both hair cells and supporting cells (Fig. 5A). This loss of cells created a large epithelial ‘wound’, that lacked any cells and likely caused disruption of the fluid barrier between endolymph and perilymph. Such epithelial wounds were evident between 5-7 days after DT treatment, but had closed by 14 days post-DT (Fig. 5B). To characterize the recovery process, we quantified cell density and epithelial repair as a function of recovery time (Fig. 5C, D). The wound perimeters were comprised of cables of filamentous actin, that were clearly labeled by phalloidin (Fig. 5E’, E”, arrows). Phalloidin labeling suggested that cells in the repaired epithelium had undergone mechanical stretching, a pattern that was consistent with epithelial closure via concentric migration of the remaining cells (Fig. 5E’’’, E’’’’). However, despite both the extensive lesion and the subsequent epithelial repair process, we did not observe nuclear translocation of YAP1 in epithelial cells at either 7- or 14-days post-DT (Fig. 5E’’’, E’’’’).

**Figure 5.**
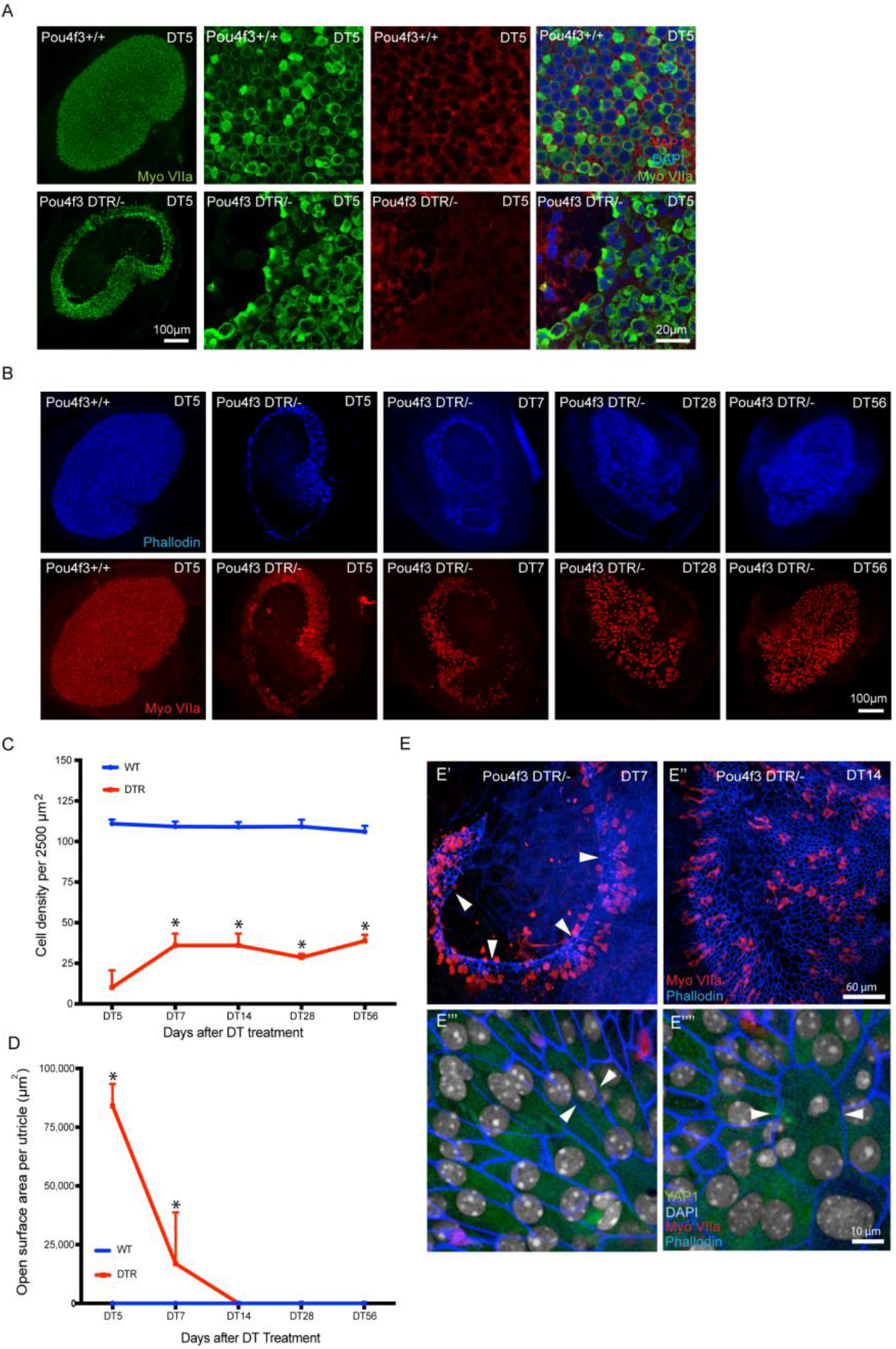
Diphtheria toxin injection creates a large epithelial ‘wound’ in utricles of neonatal Pou4f3-DTR mice. (A) Neonatal Pou4f3-DTR mice received a single 5 ng/gm injection of DT, which resulted in a large-scale loss of hair cells (myosin VIIa-green) and supporting cells, after 5 days (DT5). Such lesions were typically located in the central region of the utricle. (B) Neonatal Pou4f3-DTR mice were allowed to recover for 5, 7, 14, 28, 56 days (DT5, DT7, DT14, DT28 and DT56) after DT treatment. Wounds in the sensory epithelium (phalloidin-blue, myosin VIIa-red) were observed at 5 and 7 days after DT treatment, but were repaired by 56 days. (C, D) Quantitative analysis of cell density and epithelial repair as a function of recovery time. Data expressed as mean + SD and statistical test ANOVA was used (*p value < 0.05). N=4-6 utricle. (E) (E’) Low magnification image of the utricle from a mouse that received DT at P5 and was allowed to recover for seven days. Note that hair cells (red, myosin VIIa) and cell-cell junctions (blue, phalloidin) are missing from a large region of the epithelium. The border of the epithelial lesion is comprised of a ring of filamentous actin (arrows). (E’’) Image of utricle after 14 days recovery, showing the closure of the epithelial ‘wound’. (E’’’, E’’’’) At seven days after DT injection, the shapes of many supporting cells indicated changes in the mechanical tension within the epithelium (arrows). However, YAP1 (green) was not present in supporting cell nuclei (gray).

### Ototoxic damage to the chick utricle promotes nuclear translocation of YAP1

A final series of studies focused on the role of YAP1signaling in the chicken utricle during the initiation of regeneration after ototoxic injury. Unlike the mammalian inner ear, the auditory and vestibular organs of birds are able to quickly regenerate hair cells after acoustic trauma or ototoxicity. The molecular mechanisms that are permissive for regeneration are not known, but we hypothesized that nuclear translocation of YAP1 may be an early signal that initiates regeneration in the avian ear. We first characterized YAP1 immunoreactivity in the normal (undamaged) utricle of chickens at 2-3 weeks post-hatch. Those specimens contained ubiquitous labeling for YAP1 in the cytoplasm of supporting cells, but no YAP1 labeling in hair cells (Fig.6A). Immunolabeling for YAP1 was also present in supporting cell nuclei, but it was very rare (~1 cell/utricle; Fig. 6A, arrow). We next examined changes in YAP1 localization after aminoglycoside ototoxicity. Chicks received three injections of 1,200 mg/kg streptomycin (one/day for three days; n=5 injected chicks and 6 uninjected brood-mate controls). At 24 hr after the final injection, animals were euthanized and utricles were fixed and processed for immunohistochemical labeling. Labeling for myosin VIIa and phalloidin revealed a partial hair cell lesion in the striolar region, but very limited (or no) hair cell loss in the extrastriolar region (Fig. 6B). We quantified the numbers of cells with nuclear YAP1 immunoreactivity from three 50×50 μm striolar regions, located near the anterior, middle and posterior portions of the utricles. Such data were obtained from 9 utricles from streptomycin-treated chicks and 10 utricles from uninjected controls. The loss of hair cells in the striolar region was accompanied by increased numbers of supporting cells with YAP1-labeled nuclei (Fig 6B, arrows). In contrast, nuclear immunoreactivity for YAP1 in the extracellular region was very rare and similar to that observed in undamaged utricles (Fig 6C, D). We further examined YAP1 localization in utricles after severe hair cell lesion. For these studies, chick utricles were explanted and placed in organotypic culture, following previously described methods^29^. Utricles (n=8) were incubated for 24 hr in medium that contained 1 mM streptomycin, which results in the death >90% of the hair cell population^29^. Control utricles (n=8) were maintained in parallel, but did not receive streptomycin. All cultures were rinsed after 24 hr, fed fresh (streptomycin-free) medium and allowed to recover for 48 hr. At this point, utricles were fixed and immunolabeled for myosin VIIa and YAP1. Filamentous actin was labeled with phalloidin and cell nuclei were labeled with DAPI. Examination of myosin VIIa and phalloidin labeling revealed intact hair cells as well as a few YAP1-labeled supporting cells in control utricles (Fig. 7A, top row). However, streptomycin-treated utricles possessed severe hair cell lesions throughout the utricle, which was accompanied by nuclear immunoreactivity for YAP1 in numerous supporting cells (Fig. 7A, bottom row). Together, these data indicate that the loss of hair cells from the chick utricle leads to nuclear translocation of YAP1 protein. To determine whether YAP1 signaling was essential for the onset of regeneration, we next treated lesioned utricles with verteporfin, which blocks the association between YAP1 and TEAD cofactors and prevents DNA binding of the YAP1 complex. Utricles were placed in culture and treated 24 hr in 1 mM streptomycin. They were then rinsed and maintained for an additional 48 hr in medium that contained 1.0 μM verteporfin or 0.1% DMSO (controls, n=6 utricles/condition). Proliferating cells were labeled by addition of the BrdU to the culture medium for the final 24 hr *in vitro* (Fig. 7B). Following immunoprocessing, BrdU-labeled nuclei were quantified from three 100×100 μm regions that were distributed throughout the extrastriolar region of each utricle. Utricles treated in DMSO (controls) contained 55.1+20.7 BrdU-labeled cells/10,000 μm^2^. Treatment with 1.0 μM verteporfin reduced the level of supporting cell proliferation to 13.0+6.1 BrdU labeled cells/10,000 μm^2^ (p=0.0008) (Fig. 7C).

**Figure 6.**
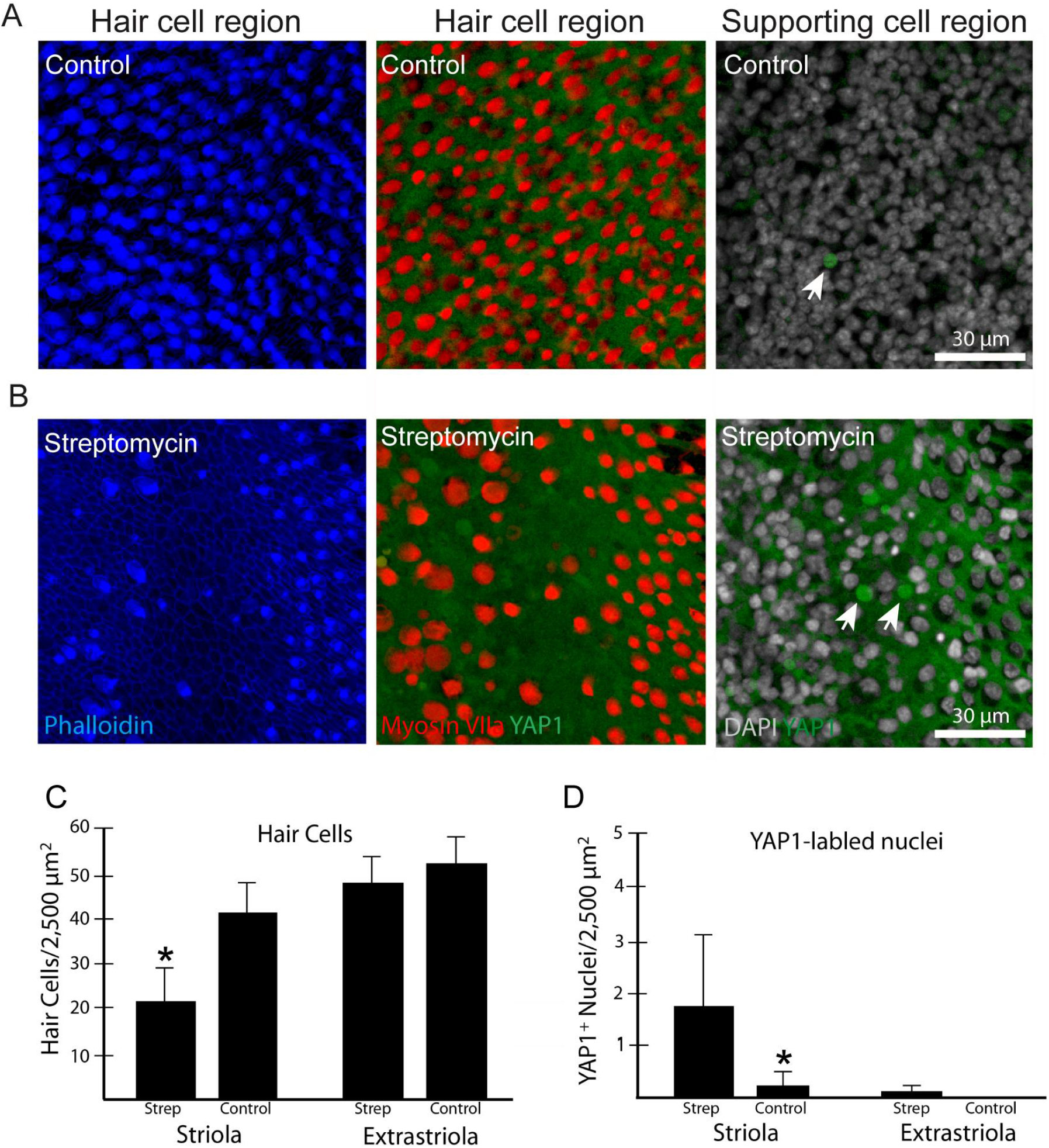
Localization of YAP1 in the chick utricle. (A) Labeling with phalloidin (blue) shows numerous stereocilia (left) and hair cells (red, labeled for myosin VIIa) surrounded by YAP1-expressing supporting cells (green) (middle). We observed a few supporting cell nuclei with nuclear YAP1 (arrow) in undamaged (control) utricles (right). (B) Effects of hair cell lesions on YAP1 localization. Chicks received 3 daily injections of streptomycin, which caused a partial loss of hair cells in the striolar region. The striolar region of injured utricles contained increased numbers of supporting cells with nuclear YAP1. (C) Quantification of hair cells in streptomycin-treated and control chick utricles. (D) Streptomycin treatment resulted in increased YAP1-labeled nuclei in the striolar region (p=0.016), but not in the extrastriolar regions.

**Figure 7.**
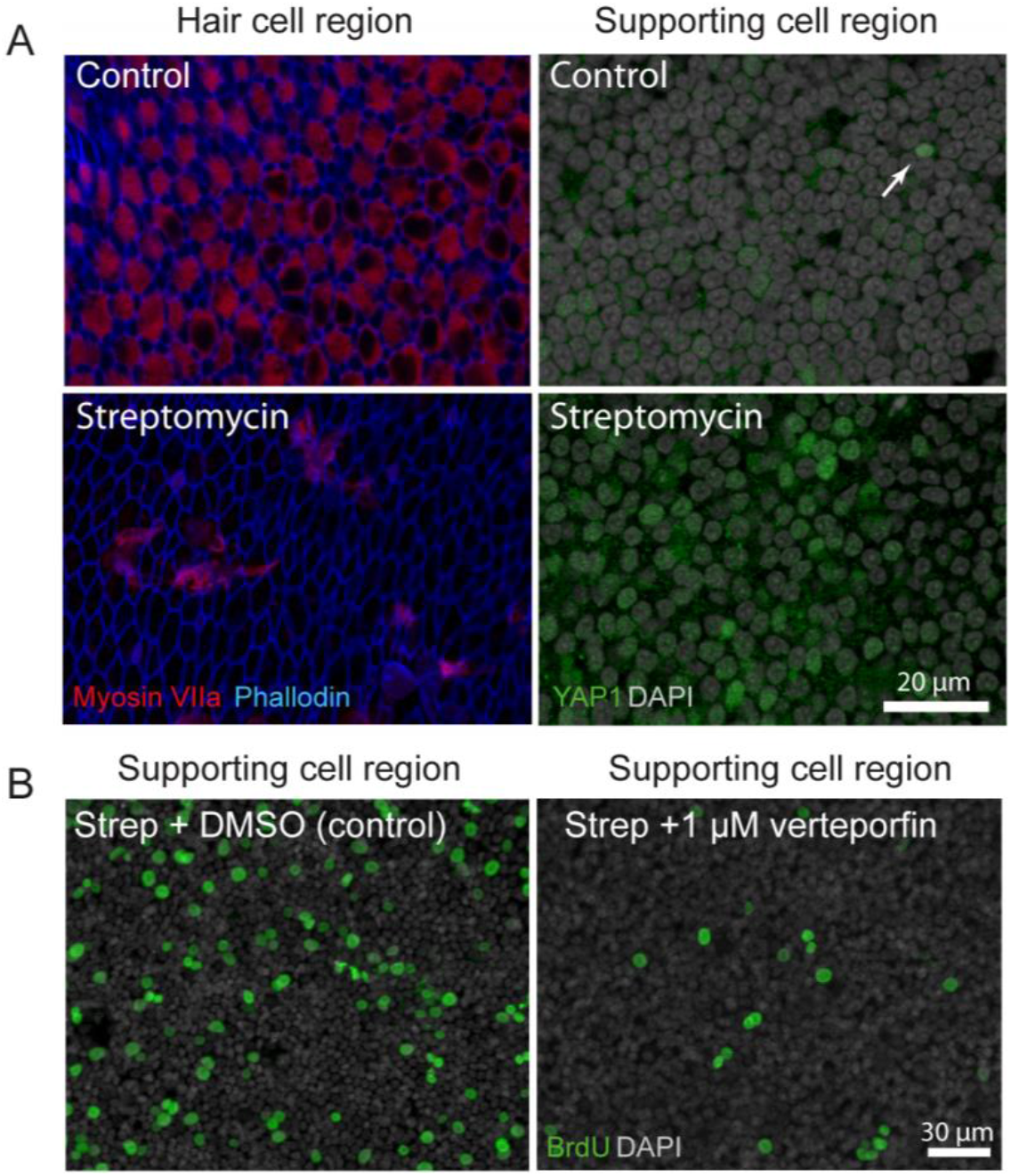
YAP1 response to ototoxic injury in organotypic cultures of the chick utricle. (A) *Left:* Utricles that were maintained in culture for 24 hr in 1 mM streptomycin show extensive loss of hair cells (blue, phalloidin), when compared to control cultures. *Right:* The hair cell lesion is accompanied by nuclear translocation of YAP1 (green) in remaining supporting cells. (B) Some utricles were allowed to recover for 48 hr after streptomycin treatment. Proliferating cells were labeled with BrdU (green), which was added to the culture medium for the final 24 hours *in vitro*. Numerous BrdU-labeled cells (green) were observed in streptomycin-injured utricles. However, addition of verteporfin caused a reduction in proliferating cells (green).

## Discussion

The objective of this study was to characterize the cellular localization of YAP1 in the utricles of mice and chicks, both in the normal ear and in response to hair cell injury. YAP1 is a transcriptional coactivator that normally resides in the cytoplasm. Under certain conditions, however, YAP1 can translocate to the nucleus, where it dimerizes with DNA-binding TEAD family proteins, and initiates changes in gene expression^5^. Nuclear translocation of YAP1 occurs after cellular injury or mechanical stress and can induce proliferation and regeneration. We observed immunoreactivity for YAP in the cytoplasm of supporting cells in utricles from both mice and chickens. Hair cell injury to the chick utricle resulted in nuclear translocation of YAP1, but we did not observe this response in the mouse utricle, even after severe epithelial injury. This suggests that the lack of YAP1 signaling following hair cell damage may be one factor that limits the regenerative ability of the mammalian ear.

One key regulator of YAP1 is the Hippo pathway, a highly-conserved signaling network that modulates cell growth and division in numerous tissues and organ systems^30^. Activation of the upstream components of the Hippo pathway determine the phosphorylation status of cytoplasmic YAP1. Phosphorylated YAP1 is targeted for degradation and does not enter the nucleus. However, activation of the upstream Hippo pathway prevents YAP1 phosphorylation, thus allowing its nuclear translocation. Once in the nucleus, YAP1 binds to TEAD family transcriptional coactivators, forming a complex that regulates gene expression. Notably, mechanical forces exerted on cells can also influence YAP1 phosphorylation, although it is not clear whether such forces act via the Hippo pathway or by other signaling mechanisms^31^.

Although we did not observe nuclear translocation of YAP1 in the injured mouse utricle, we did observe nuclear entry of YAP1 after utricles were removed from mice and placed in culture. This response was rapid and transient. Nuclear immunoreactivity for YAP1 was noted after two hours *in vitro*, but was not present after 24 hr of culture. The processes of dissection and placement in organotypic culture are likely to dramatically alter the mechanical environment of the utricle, and the observed changes in YAP1 are consistent with the notion that mechanical forces influence YAP1 localization^6,32–35^. It is possible that hair cell loss or epithelial wounding may cause local changes in cellular tension, but that large-scale mechanical disruptions are required for activation of YAP1 signaling. Our data from the cultured explants also indicate that YAP1 translocation occurs at very short intervals after changes in epithelial mechanics (i.e., within 2 hr of dissection). It is possible that YAP1 is also translocated to the nuclei of supporting cells at similarly short times after epithelial injury, but that its expression once again becomes cytoplasmic within 24 hr after injury. The temporal resolution of our methods does not permit us to resolve this issue, i.e., lesions created by DT injection in Pou4f3-huDTR mice are not synchronized and require 3-7 days to appear^27–28^.

We also observed cytoplasmic immunoreactivity for YAP1 in a subset of hair cells in the developing utricle. Such cells were only observed during the first postnatal week. They also possessed Sox2-labeled nuclei (a marker of type II identity^36, 37^) and were generally concentrated in the lateral portion of the sensory epithelium. Prior studies have shown that most type II hair cells of the mouse utricle differentiate during the first postnatal week and are disproportionally added to the lateral region of the sensory epithelium^38–39^. It is likely that the YAP1-labeled hair cells had recently differentiated from YAP1-expressing precursors and were undergoing differentiation as type II hair cells. This suggestion is consistent with single cell RNA seq data obtained from utricles of newborn mice^40^, which indicate that YAP1 is expressed in immature hair cells (data available at umgear.org). However, interactions between YAP1 and Sox2 are involved in the maintenance of stemness and fate determination in different types of stem cells^41–44^, and it is possible that YAP1 and Sox2 interactions may serve some unidentified role in the development of the utricle.

One unexpected observation reported here is the presence of a large epithelial ‘wound’ in the utricles of neonatal Pou4f3-huDTR mice after a single 5 ng/gm injection of DT. This treatment reliably caused extensive loss of both hair cells and supporting cells, resulting in a large opening in the sensory epithelium. Such lesions were not observed in the cristae of the semicircular canals or in the cochlea; in those sensory organs, DT injection caused a selective loss of hair cells, a pattern which resembled that reported after DT treatment of mature Pou4f3-huDTR mice^18–27–28^ (Warchol, unpublished data). The reason for the creation of the epithelial wound is not clear. It is possible that some supporting cells in the neonatal utricle transiently express Pou4f3, leading to co-expression of the DT receptor in those cells. Also, the E-cadherin-mediated cellular junctions in the neonatal utricle are not mature at P5^45^, so the epithelium may not be able to maintain integrity after a high level of DT-mediated hair cell death. We further found that these lesions closed spontaneously within seven days, similar to the pattern of closure observed after *in vitro* puncture wounds in the utricles of embryonic mice^46^. Such large epithelial wounds are likely to have caused considerable disruption in the mechanical environment experienced by the remaining cells. Further changes in cellular tension would also occur during the process of wound closure. Notably, however, none of these changes was sufficient to cause nuclear translocation of YAP1. Such translocation has been observed following lesioning of the sensory epithelium and stromal tissue of the neonatal mouse utricle^6^, but the types of lesions employed in these two studies - DT-mediated epithelial wounding vs. tissue injury caused by a fine needle - are significantly different. The needle-induced lesion used by Gnedeva et al., (2017) may have applied considerably more mechanical force on the sensory epithelium than was created by our DT lesion.

Finally, our data suggest important functional differences in YAP1 signaling in the utricles of birds vs. mammals. Cytoplasmic YAP1 was observed in all supporting cells of the chick utricle. In addition, nuclear YAP1 was observed in supporting cell nuclei after hair cell lesions created either *in vitro* or *in vivo*, and disrupting YAP1 signaling by treatment with verteporfin resulted in a reduction in regenerative proliferation. These observations suggest that YAP1 may serve a role in the initiation of regeneration in the avian inner ear. As noted above, we observed no changes in YAP1 localization after hair cell injury in the utricles of neonatal or mature mice, so it is possible that differences in YAP1 signaling may partially explain the differing regenerative abilities of the avian vs. mammalian inner ear. They also raise the possibility that activation of Hippo/YAP1 signaling in mammals may lead to some degree of hair cell regeneration.

## Methods

### Animals

Studies used mice of both sexes on C57BL/6 or CD1 backgrounds. Some studies also used Pou4f3-huDTR transgenic mice, in which the human form of diphtheria toxin receptor (hu-DTR) gene is expressed under the control of the Pou4f3 transcription factor promoter^18, 27, 47^. Chickens were hatched from fertile eggs (Charles River SPAFAS) and maintained in heated brooders until used in experiments. Mice and chickens were housed in the animal facilities of Washington University in Saint Louis, and were maintained on a 12-hr/day-night light cycle with open access to food and water. All experimental protocols involving animals were approved by the Animal Studies Committee of the Washington University School of Medicine, in Saint Louis, MO.

#### Experiments with mice

##### Genotyping

Genotyping protocol for identification of *Pou4f3*^DTR/+^ and *Pou4f3^+/+^* was similar to Tong et al, (2015). Briefly, DNA was extracted from tails using ethanol precipitation. PCR was used to amplify the targeted allele (Quick-Load^®^ *Taq* 2X Master Mix, New England Biolabs Inc), using the following primers (at 0.4μM): *Pou4f3* (WT) Forward 5’ CAC TTG GAG CGC GGA GAG CTA G; *Pou4f3* (mutant) Reverse 5’ CCG ACG GCA GCA GCT TCA TGG TC. PCR was performed using the following reaction conditions: 95° C for 5 min; 95° C for 30 s, 59° C for 30 s, 72° C for 1 min, 30 cycles; 72° C for 7 min; 4° C infinity. PCR products were run on 1.5-2% agarose gel containing 1μl/ml SYBR safe DNA gel stain (expected band~ 150 bps) (ThermoFisher).

##### Hair cell ablation

Mice received a single dose of Diphtheria toxin (DT, Sigma), which was administered intramuscularly (i.m., 5 ng/gm) in the thigh region of the hind leg of P5 mice and intraperitoneally (i.p., 25 ng/gm) in 4-6-week adult mice. Using identical methods, DT was also administered to wild type (WT) littermates, which served as controls. Mice were allowed to survive for 5, 7, 14, 28, or 56 days after DT injections.

##### Utricle explant culture

Mice were euthanized at P15 or P28 and temporal bones were removed and placed in tissue culture medium under sterile condition. Utricles were isolated and otoconia were gently removed from the surface using fine forceps. Utricles were cultured free-floating and/or attached to Matrigel-coated surfaces in 1 cm diameter wells (MatTek). Each well contained 100 μl of Medium 199 with Earle’s salts, 2200 mg/L sodium bicarbonate, 0.69 mm l-glutamine, 25 mm HEPES (Gibco), supplemented with 10% FBS and 10 μg/ml Ciprofloxacin. Utricles were cultured at 37°C in a 5% CO2/95% air environment.

##### Immunohistochemistry

For *in vivo* samples, mice were euthanized with Fatal Plus and isolated temporal bones were fixed with 4% paraformaldehyde (PFA) in 0.1M phosphate buffer (PB) overnight at 4° C. Cultured utricles were fixed for 1-2 hr with 4% paraformaldehyde in PB at room temperature. After fixation, utricles were washed 3x (5 min each) in PBS and then processed for whole mount immunohistochemistry. Rabbit polyclonal anti-Myosin VI antibody (catalog #25–6791, Proteus BioSciences, 1:500) was used to label hair cells, Goat polyclonal anti-Sox2 antibody (catalog # sc-17319, Santa Cruz Biotechnology, 1:100) was used to label supporting cells, and two different YAP1 antibodies were used to characterize YAP1 expression patterns: mouse monoclonal anti-YAP1 antibody (catalog # sc-101199,1:50) and rabbit monoclonal anti-YAP1 antibody (catalog # 14074S, Cell Signaling,1:100). To prevent nonspecific binding of the antibodies, samples were incubated in a blocking solution consisting of 5% normal horse serum/0.2% Triton X-100 in PBS for 1 hr at room temperature. Samples were then incubated overnight in primary antibodies prepared in PBS with 2% horse serum and 0.2% Triton X-100 at 4° C. Samples were then rinsed 3x (5 min each) in PBS and incubated for 2-3 hr in secondary antibodies (conjugated to Alexa-488, Alexa-568, and Alexa 647, Life Technologies,1:500) at room temperature. All secondaries were prepared in PBS with 2% horse serum and 0.2% Triton X-100. Filamentous actin was labeled with Alexa Fluor^®^647 Phalloidin and Alexa Fluor^®^488 (Invitrogen, catalog #A22287, and #A12379, 1:200), and cell nuclei were labeled with DAPI (catalog #D9542, Sigma-Aldrich, 1 μg/ml). All samples were rinsed 3x (5 min each) in PBS. Samples were mounted in glycerol: PBS (9:1) solution and coverslipped on glass slides.

#### Studies involving chickens

Hatchling chicks were housed in Washington University animal facilities as described above. Studies conducted *in vivo* used chickens at ~4-week post-hatch. Chickens received injections of streptomycin sulfate (1,200 mg/kg, i.m.) once/day for three consecutive days. Chickens were allowed to recover for 24 hr after the last injection and were then euthanized by CO_2_ inhalation. Utricles were quickly removed and fixed for 30 min in 4% paraformaldehyde (in 0.1 M PB). Specimens were rinsed 3x and processed for immunohistochemical labeling, using methods described above. Controls consisted of age-matched (clutch-mate) chickens that did not receive streptomycin.

Organotypic cultures of chick utricles were prepared following previously described methods^29^. Briefly, chicks (10-20 days post-hatch) were euthanized via CO_2_ inhalation and quickly decapitated. The lower jaw and skin covering the head were removed and heads were immersed for 5-10 min in 70% EtOH. All subsequent work was conducted under aseptic conditions. Utricles were removed from temporal bones and transferred to chilled Medium-199 (with Hanks salts and 25 mM HEPES; Thermo-Fisher). The otoconia were removed and isolated sensory organs were placed in 1 cm culture wells (Mat Tek, Ashland MA) that contained 100 μl of Medium-199 (with Earles salts, 2,200 mg/L sodium bicarbonate, 0.69 mM L-glutamine and 25 mM HEPES) supplemented with 1% fetal bovine serum (FBS). Some utricles also received streptomycin, for a final concentration of 1 mM. Utricles were incubated at 37° C a humid 5% CO_2_/95%air environment. After 24 hr *in vitro*, specimens were rinsed 3x with fresh medium and given 100 μl of Medium-199/1%FBS, that also contained 1.0 μM verteporin (Sigma) or 0.1% DMSO (vehicle). Cultures were maintained in these media for an additional 48 hr and BrdU (3 μg/ml) was added for the final 24 hr *in vitro*. Specimens were fixed for 30 min in 4% PFA and processed for immunocytochemical labeling of BrdU (protocol in Slattery and Warchol, 2010). Nuclei were counterstained with DAPI. Specimens were visualized as wholemounts and confocal images were obtained from three 100×100 μm regions in the extrastriolar portion of each utricle.

##### Cellular imaging and analyses

Fluorescent images were obtained using a Zeiss LSM 700 confocal microscope. For all specimens, Z-series images were obtained at 10× (~4.5-micron z-step size), 20× (1 or 2 -micron z-step size), or 63× (0.5 or 1.0-micron z-step size) objectives. Images were processed and analyzed using Volocity 3D image analysis software (version 6.3, PerkinElmer) and Fiji (ImageJ2.0) (National Institutes of Health) and Adobe illustrator CS5.1.

##### Hair cell and supporting cell counts

Cell quantification was performed from 63x images using Fiji software (ImageJ2.0, National Institutes of Health). For all cell counts, lateral extra striolar, striolar and medial extra striolar region were selected per utricle samples. The Cell Counter plugin was used for all cell counts. For hair cell counts, a grid of 1,000 μm^2^ or 2,500 μm^2^ was applied to the Z-stack images. Hair cells were identified by strong cytoplasmic immunolabeling for myosin Vlla. All hair cells were manually counted within the designed area of complete z-stack. For supporting cell counts, a grid of 10,000 μm^2^ was applied to the Z-stack images. Supporting cells were identified by strong nuclear immunolabeling for Sox-2 protein. All supporting cells were manually counted within the designed area of complete z-stack.

##### Statistical analysis

All the data analysis and statistics were carried out using GraphPad Prism version 6.0d. Data are presented as mean ± SD. Student’s t-tests or analyses of variance (ANOVA) followed by Tukey’s or Bonferroni’s post hoc tests were applied, as appropriate. Results were considered statistically significant when p < 0.05.

